# Knockout of *fatty acid elongase1* homeoalleles in amphidiploid *Brassica juncea* leads to undetectable erucic acid in seed oil

**DOI:** 10.1101/2024.10.09.617390

**Authors:** Neelesh Patra, Guy C. Barker, Mrinal K. Maiti

**Author notes:** **Corresponding author** ^1^Mrinal K. Maiti.

## Abstract

Indian mustard (*Brassica juncea* L.) is a major oilseed crop with significant economic and nutritional importance within the Indian subcontinent. While its seed oil offers valuable dietary benefits, including a balanced ratio of human essential fatty acids, the traditional high oil-yielding varieties contain an elevated level of erucic acid (EA, C22:1) that is associated with adverse health effects. Therefore, developing low erucic acid (LEA) mustard cultivars is crucial for broader utilization and consumer safety. In this study, we employed CRISPR/Cas9 genome editing tools to disrupt the *fatty acid elongase1* (*FAE1*) gene that encodes a key enzyme in EA biosynthesis in two high erucic acid (HEA) *B. juncea* cultivars, PCR7 and JD6. Targeted knockout (KO) of *BjFAE1* homeoalleles (*BjFAE1.1* and *BjFAE1.2*) in this amphidiploid plant using CRISPR/Cas9 constructs, each carrying two guide RNAs led to monoallelic and biallelic mutations. Biallelic KO lines showed a near-complete elimination of EA (<0.5% in PCR7, undetectable in JD6) with a significant increase in nutritionally beneficial oleic acid (∼30% in PCR7, ∼35% in JD6), while the content of essential fatty acids also increased significantly, suggesting rerouting of carbon flux from EA biosynthesis. Importantly, these LEA lines retained key agronomic traits like plant seed yield and oil content, matching the productivity of the control elite cultivars. Our findings underscore the effectiveness of CRISPR/Cas9 technology for editing *B. juncea* genome, producing LEA seed oil lines with improved nutritional quality and thus expanding the applications of this important oilseed crop.

## Introduction

Most plant species biosynthesize and accumulate storage compounds in their seeds as reservoirs of carbon and energy, crucially needed for successful propagation. These storage materials, primarily carbohydrates (often starch), lipids (predominantly triacylglycerols), and specialized storage proteins, serve the sole purpose of seedling development until photosynthetic energy independence is achieved (Taiz and Zeiger, 2010). Like other seed-stored biomolecules, plant lipids or oils, with their diverse fatty acids, hold immense economic value as a sustainable source of human nutrition, animal feed, and feedstocks for various food and non-food industrial applications. The proportion of the different fatty acid (FA) molecules that vary in chain length and degree of saturation, dictate the physicochemical properties and suitability of the oil for specific end usage (Gunstone, 2011).

Among oilseeds, rapeseed-mustard (*Brassica* sp.) holds a prominent position globally, ranking third in terms of production volume, trailing only oil palm and soybean (Foreign Agricultural Service/USDA, 2024). In India, rapeseed-mustard remains a significant contributor to nation’s agricultural economy, accounting for 31% of the country’s oilseed production (Government of India, 2024). Indian mustard (*Brassica juncea* L.), a natural amphidiploid or allotetraploid (AABB) formed by the hybridization of *B. rapa* (AA) and *B. nigra* (BB) genomes, displays immense value as a key oilseed crop in Indian subcontinent exhibiting wide climatic adaptability across the countries (Kang et al., 2021). Seed oil of Indian mustard presents a duality for both human and animal nutrition. Although it contains a very low amount of saturated FAs, moderate amount of beneficial FAs such as ω-9 oleic acid (C18:1), and human essential FAs [i.e., ω-6 linoleic acid (C18:2), and ω-3 linolenic acid (C18:3)], it also has a high level of ω-9 erucic acid (C22:1), a monounsaturated very-long-chain fatty acid (VLCFA) (Singh et al., 2014). Generally, VLCFAs with 20 carbons and more in chain length, are essential building blocks for various plant structures and biomolecules, including cell membranes, protective waxes on the outer layer (cuticle), and energy-rich storage lipids within seeds (Bach and Faure, 2010). Though valuable for industrial usage in the synthesis of nylon, plastics, and lubricants (Przybylski, 2011), the high erucic acid (EA) content in food and feedstock can have adverse health implications. Accumulating evidence suggests that high EA consumption may have adverse effects on human health, particularly in relation to cardiovascular function, such as the development of myocardial lipidosis (accumulation of fat within heart cells) and increased blood cholesterol level, as documented in some mammalian models (Abdellatif and Vles, 1971; Mortuza et al., 2005). Additionally, low EA mustard oil has the potential as a sustainable feedstock for biofuel production (Knothe, 2010).

EA is mainly found in the seeds of many *Brassicaceae* family plant species. The biosynthesis of EA involves a four-step elongation process wherein the *Fatty Acid Elongase 1* (*FAE1*) gene product plays a central role. The 3-ketoacyl-CoA synthase (KCS), encoded by *FAE1* catalyzes the rate-limiting and substrate-specific initial condensation step for extending the fatty acyl chain length from C18:0- or C18:1-CoA (Coenzyme A) to initiate VLCFA synthesis (Millar and Kunst, 1997; Li-Beisson et al., 2010). The diploid *Brassica* species (*B. rapa* and *B. nigra*) possesses a single copy of *FAE1* gene, whereas the amphidiploid species (*B. napus* and *B. juncea*) have two copies, viz., *FAE1.1* and *FAE1.2*, contributing additively to seed EA accumulation (Gill et al., 2021). Simultaneous disruption of *FAE1* homeologs via CRISPR/Cas9 provides a promising strategy to effectively block EA biosynthesis, enhancing the oil’s overall nutritional value.

Traditional breeding efforts paved the way for developing low erucic acid (LEA) cultivars in some *Brassicaceae* family plants. Notably, Canadian rapeseed breeders pioneered the creation of LEA varieties of *B. napus* and *B. rapa* (Downey and Craig, 1964). When combined with the genetic trait of low glucosinolate content, these LEA varieties led to the development of "canola" oil – a registered trademark denoting rapeseed-mustard containing less than 2% EA in seed oil and less than 30 µmol of glucosinolates per gram of oil-free meal (FCC standard for canola oil, 1996). Similar success in *B. juncea* followed the discovery of LEA (<2%) genotypes (Zem 1 and Zem 2) in Australia (Kirk and Oram, 1981). However, the double recessive nature of the LEA trait and the possibility of linkage drag make it hard to use traditional breeding methods to create LEA elite Indian mustard cultivars that do well in a wide range of agroclimatic zones across the Indian subcontinent (Gill et al., 2021).

Currently, in India, only a limited number of canola-quality mustard varieties and hybrids are being cultivated due to undesirable agronomic features such as delayed maturation, smaller seed size, and lower seed oil content in those lines than elite cultivars (Banga et al., 2015). In contrast to the global trend towards LEA cultivars, Indian mustard varieties with 40–50% seed EA content remain predominant (>90%) among commercially cultivated rapeseed-mustard varieties in India and other countries of Indian subcontinent like Pakistan and Bangladesh, due to their superior oil yields (Iqbal et al., 2008; Mortuza et al., 2005). The development of LEA cultivars with optimal agronomic performance remains a priority to broaden the utilization of Indian mustard (*B. juncea*) for safer human consumption and expand its industrial applications.

The advent of precise genome editing tools like CRISPR/Cas9 offers new opportunities for overcoming these limitations and accelerating the development of LEA cultivars with desirable agronomic traits. Recent research in *B. napus* demonstrated the potential of CRISPR/Cas9 in creating low EA germplasms by targeting both copies of the *FAE1* gene (BnaA08.FAE1 and BnaC03.FAE1), resulting in near zero EA levels with no significant impact on key agronomic traits in three edited germplasms (Liu et al., 2022). In present study, the CRISPR/Cas9 tool was employed in complex amphidiploid genome in *B. juncea* to achieve targeted disruption of the *FAE1* genes at multiple loci.

Successfully employing CRISPR/Cas9 technology to disrupt *BjFAE1* homeoalleles (*BjFAE1.1* and *BjFAE1.2*) in two elite cultivars, PCR7 and JD6, of *B. juncea* remarkably reduced the EA level in seed oil. In line with expectations, these genome-edited lines exhibit significantly increased content of nutritionally desirable FAs, such as oleic acid, linoleic acid and linolenic acid in seed oil. Importantly, these CRISPR/Cas9 knockout lines demonstrate plant growth parameters (vegetative and reproductive) and oil content comparable to unedited control cultivars.

## Results

### Sequence analysis and expression profiling of *fatty acid elongase1* genes of two *B. juncea* cultivars accumulating high EA in seed oil

Five Indian mustard (*B. juncea*) cultivars (NPJ-112, NPJ-113, NPJ-124, PCR7, and JD6) were examined for seed oil content by gravimetric quantification and FA compositions using gas chromatography-mass spectrometry (GC-MS). Two cultivars, PCR7 (Rajat) and JD6 (Pusa Mahak) were selected for further study due to their higher seed oil content of ∼40% and ∼45%, respectively (**Figure 1a, Table S1**) and higher EA content of ∼39% and ∼45%, respectively, in seed oil (**Figure 1a, 1b, Table S2**) compared to the other cultivars.

**Figure 1.**
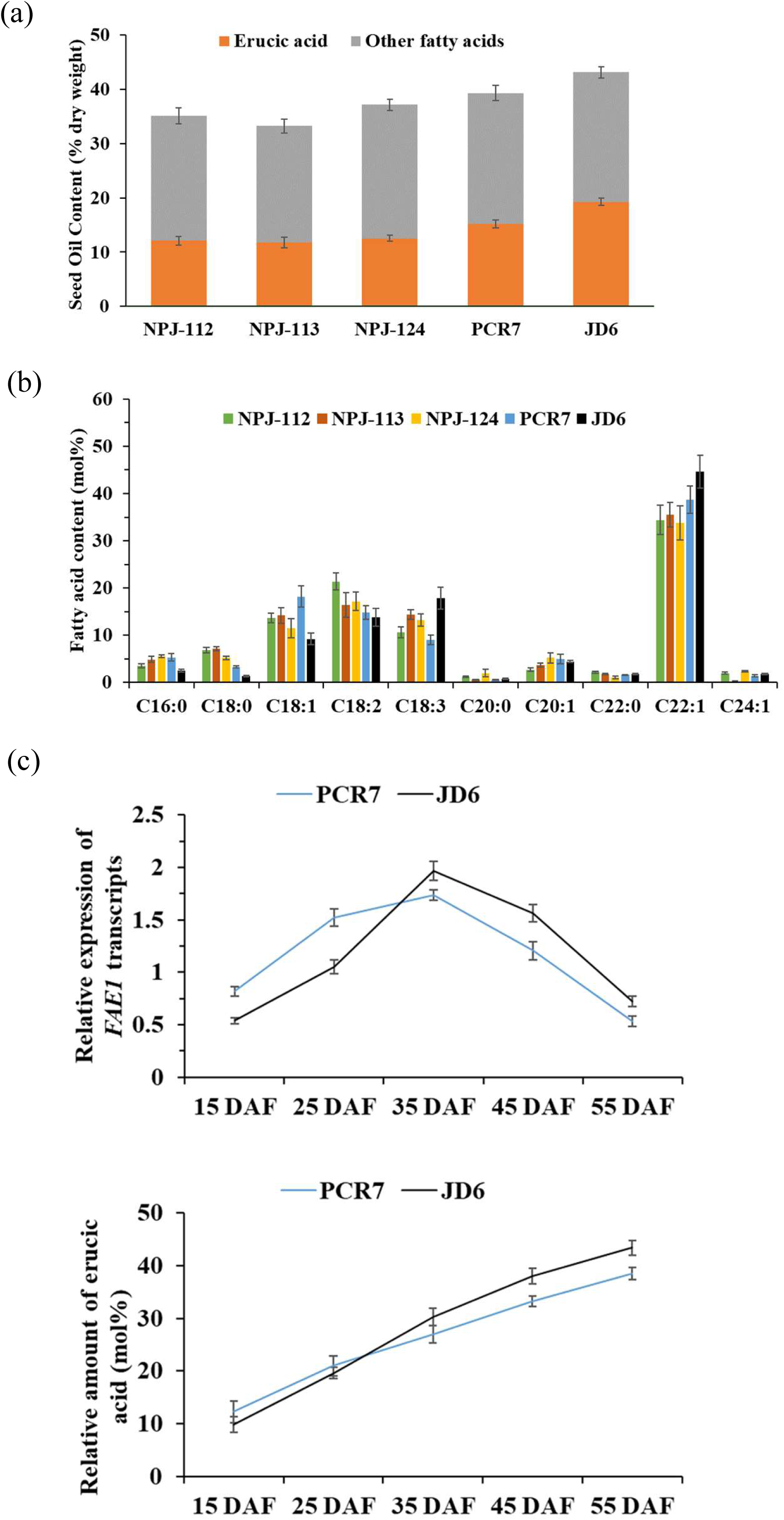
Selection of *B. juncea* cultivars with a high content of seed oil and erucic acid (EA), and expression profiling of *BjFAE1* gene in two selected cultivars. (**a**) Seed oil content of five Indian mustard cultivars viz. NPJ-112, NPJ-113, NPJ-124, PCR7 and JD6 along with the EA level in their seed oil. Values represent mean ± standard error (SE) of five to seven independent experiments. (**b**) Seed fatty acid (FA) composition (mol%, mean ± SE, n=5) of the five cultivars mentioned in (a). (**c**) *BjFAE1* transcript levels (relative to elongation factor 1-alpha (EF1α)) and EA content (mol% of total FAs) in developing seeds of PCR7 and JD6 cultivars at 15, 25, 35, 45, and 55 days after flowering (DAF). Values represent mean ± SE (n=3).

Both copies of *FAE1* genes, *BjFAE1.1* and *BjFAE1.2*, responsible for regulating the level of EA in seed oil were cloned and sequenced from the genomic DNA and seed-specific cDNA from PCR7 and JD6 cultivars. *In silico* analysis validated the presence of two highly similar gene copies (homeoalleles) in the amphidiploid *B. juncea* genome and the intron-less nature of the gene. Investigation of the expression pattern of these genes through the use of real-time PCR confirmed the seed-specific expression of *FAE1* transcripts, with peak expression occurring during mid to late seed development, coinciding with the incremental EA accumulation in developing seed tissues (**Figure 1c, Table S3**).

### Preparation of CRISPR/Cas9 constructs with two guide RNA expression cassette targeting *FAE1* homeoalleles

In order to reduce the content of nutritionally undesirable EA in seed oil of PCR7 and JD6 cultivars, CRISPR/Cas9 technology was employed for simultaneous disruption of *BjFAE1* homeoalleles (homoeologs) that encode the FAE1 enzyme crucial for EA biosynthesis (**Figure 2a**). To create functional knockout of the *FAE1* gene, four suitable single guide RNAs (sgRNA1 - 4) (**Table S4**) targeting conserved regions of *BjFAE1.1* (denoted as E1) and *BjFAE1.2* (denoted as E2) coding sequence (**Figure 2b**) were designed through *in silico* analysis considering all the factors necessary for maximal activity of sgRNA-Cas9 complex. Two final chimeric constructs, viz., pBjFAE13 and pBjFAE24, were prepared based upon *Agrobacterium tumefaciens* Ti-plasmid (**Figure 2c**), both carrying SpCas9 gene controlled by CsVMV promoter, hygromycin resistance gene controlled by CaMV35S promoter, and a combination of two sgRNAs individually driven by the AtU6 promoter. Thus, pBjFAE13 harboured sgRNA1 and sgRNA3, likewise pBjFAE24 harboured sgRNA2 and sgRNA4 for simultaneous targeting a pair of genomic regions of *FAE1* coding sequences to introduce insertion or deletion of a few nucleotides at Cas9 cleavage site leading to functional inactivation of E1 and E2 alleles by frameshift mutation. Deletion efficiency of each construct was checked by transient protoplast transformation before creation of stable transgenic lines.

**Figure 2.**
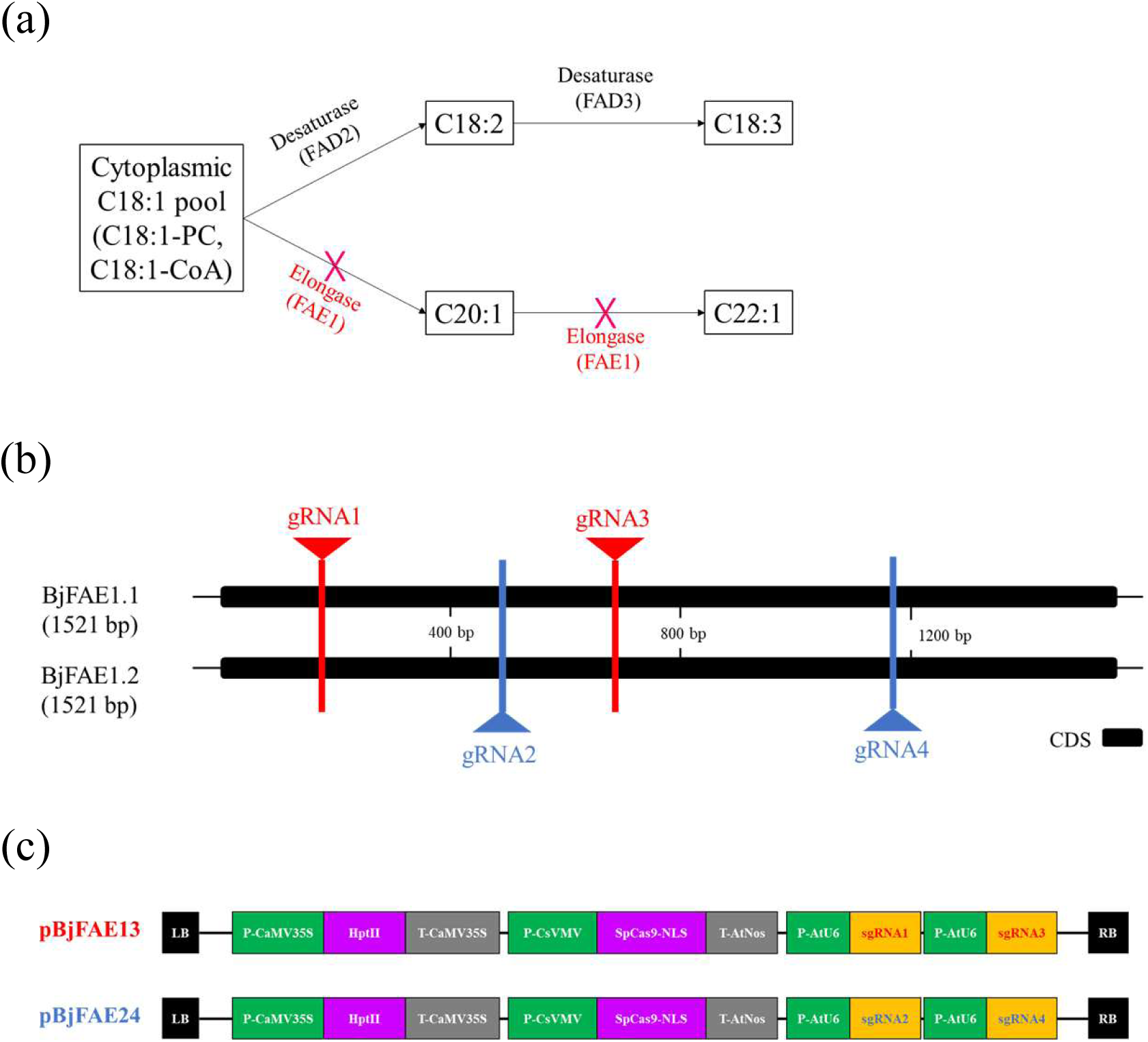
CRISPR/Cas9 constructs each carrying the expression cassette of two sgRNAs simultaneously for targeted knockout of *BjFAE1* homeoalleles in amphidiploid *B. juncea*. (**a**) Illustrative biochemical pathways for desaturation and elongation of FAs in seed tissues of *B. juncea*. Red cross indicates mutation of *FAE1* genes encoding fatty acid elongase1 enzyme to block the biosynthesis of EA. (**b**) Genomic location of the target sites of four guide sequences designed for disrupting *BjFAE1* coding sequence of two homeoalleles *BjFAE1.1* and *BjFAE1.2*. (**c**) T-DNA map showing *Agrobacterium tumefaciens*-based transformation constructs, viz., pBjFAE13 and pBjFAE24 used for targeting the *BjFAE1* homeoalleles in two *B. juncea* cultivars, PCR7 and JD6.

### Genetic transformation of *B. juncea* and development of *BjFAE1*-knockout lines

The knockout constructs, pBjFAE13 and pBjFAE24, were introduced separately into *A. tumefaciens* strain LBA4404 for subsequent transformation of hypocotyl explants of *B. juncea* cultivars PCR7 and JD6 according to the reported protocol (Bhattacharya et al., 2015) with minor modifications. After *Agrobacterium* infection, the plantlets regenerated from hypocotyls were screened on 30 mg/L of hygromycin-supplemented medium (**Figure 3a**) for ten consecutive culture passages (the first six culture passages on shoot induction medium and the final four culture passages on root induction medium). Surviving plantlets (**Figure 3b**) were transferred to greenhouse (**Figure 3c**). Genomic integration of the T-DNA segment of CRISPR/Cas9 constructs was confirmed by hygromycin- and Cas9-specific PCRs (**Figure 3d**). Based on PCR screening results, 27 out of 72 T0 plantlets of PCR7 cultivar and 32 out of 86 T0 plantlets of JD6 cultivar which proved positive were grown till maturity, and seeds were harvested separately from each line. Further molecular confirmations of T0 lines were carried out by high resolution melting (HRM) analysis (**Figure 3e**) of ∼200 bp PCR amplified fragment encompassing a probable Cas9 cut site, followed by Sanger sequencing and Inference of CRISPR Edits (ICE) analysis (**Table S5**) using the obtained sequencing data.

**Figure 3.**
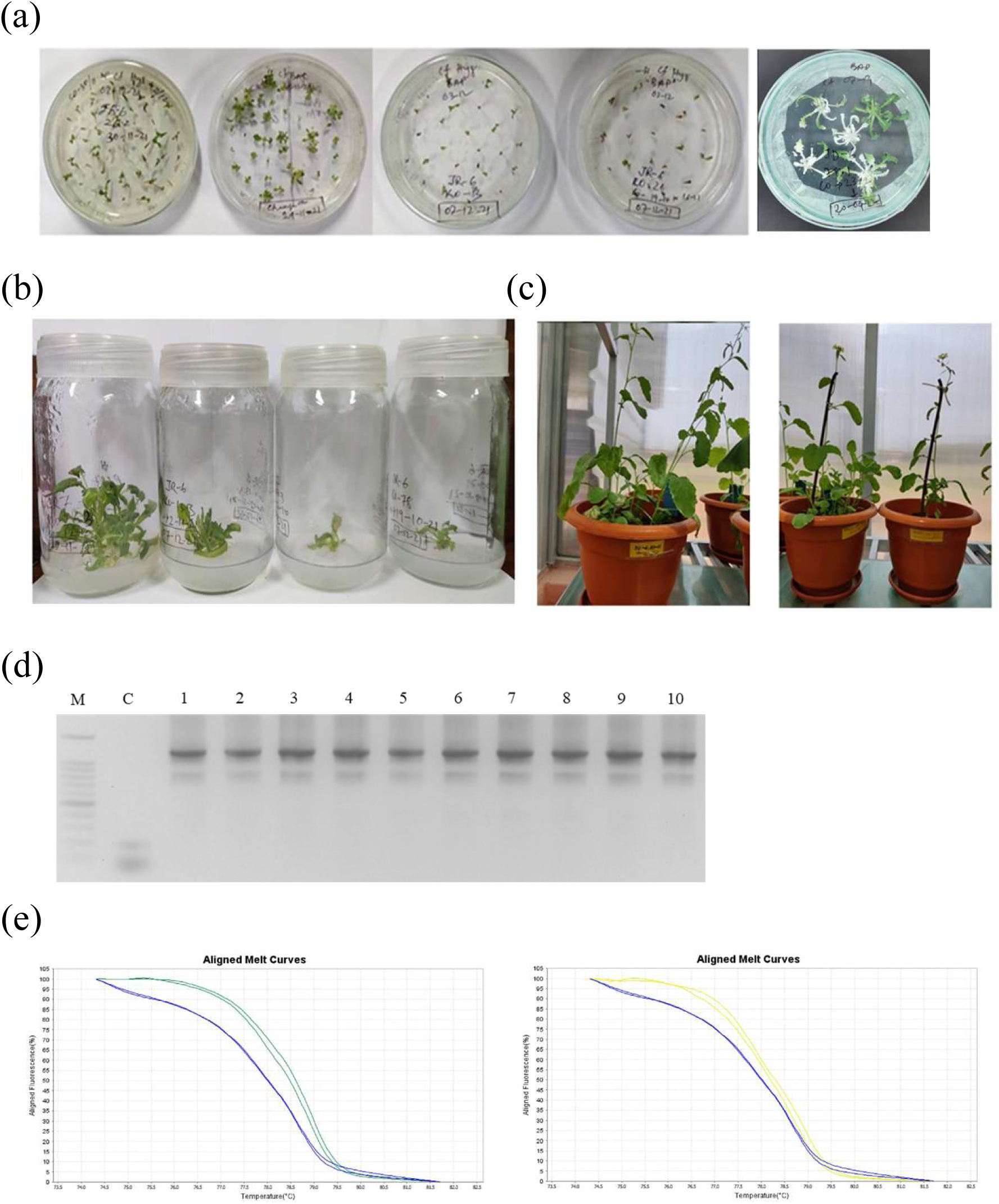
Development of *BjFAE1* knockout T0 mustard lines using CRISPR/Cas9 constructs. (**a**) *In vitro* regeneration of plantlets after *B. juncea* hypocotyl transformation by *A. tumefaciens* LBA4404 strain harboring either pBjFAE13 or pBjFAE24 KO construct. (**b**) A few hygromycin selected plantlets in root induction medium. (**c**) Hygromycin resistant and Cas9 gene PCR-positive T0 plants growing in controlled greenhouse till seed collection. (**d**) PCR amplification of transgene-specific ∼1250 bp amplicon from untransformed and potential KO transgenic T0 plants using Cas9 gene specific forward primer and T-DNA right border specific reverse primer. (**e**) HRM analysis of putative KO lines of T0 generation.

### Inheritance pattern of the transgene in T1 progenies and characterization of stable *FAE1* knockout lines in T2 progenies

We investigated inheritance patterns of the T-DNA segment from pBjFAE13 and pBjFAE24 knockout (KO) constructs in transgenic progenies of each cultivar separately. The T-DNA segment of these KO constructs carries linked genes for hygromycin resistance (Hyg^R^), Cas9 and two sgRNA expression cassettes. T0 transgenic plants from multiple independent transformation events were self-fertilized in a controlled greenhouse. For each cultivar, twenty T0 lines were randomly chosen. Seeds were harvested from each T0 lines separately, and the progenies of each line were analyzed for transgene inheritance and segregation with respect to Hyg^R^ phenotype and Cas9 gene integration.

All harvested seeds from T0 lines were germinated separately on Murashige and Skoog medium with hygromycin. Hygromycin exposure caused death for all the seedlings from untransformed controls and up to 60% seedlings of T1 lines. PCR with Cas9-specific primers confirmed the presence of Cas9 gene in all the randomly selected subset of 5 to 8 hygromycin-resistant seedlings per T0 line. Overall, progeny from 11 PCR7 and 14 JD6 transgenic lines exhibited an approximate 3:1 Mendelian segregation ratio for Hyg^R^ phenotype, suggesting a single-copy integration of the T-DNA segment /transgene (**Table S6**). Progenies from the remaining 9 PCR7 and 6 JD6 transgenic lines displayed ∼15:1 or other segregation ratios, indicating two or more transgene integration sites (**Table S6**).

T2 progenies were generated by self-pollination from selected Hyg^R^ and Cas9 positive seedlings from each single-copy transgene integration T1 line and were grown till maturity. T7 endonuclease I surveyor assay (**Figure 4a**) was used to screen for insertion-deletion (indel) mutations in the target regions of the *FAE1* gene in T2 progenies, and a few biallelic mutant lines were confirmed by Sanger sequencing (**Figure. 4b**). Sequence analysis also revealed some T2 progeny plants with single allele (either E1 or E2) mutations. Analysis of the mutated sequences revealed that CRISPR/Cas9 editing generated a variety of indels at the target site, ranging from 60 base pair additions to deletions of 40 base pairs (**Figure 4b**). To verify the unintended mutations in T2 lines caused by the sgRNAs, predicted off-target sites were sequenced in three randomly selected e1e2 plants per cultivar. No unintended mutations were detected.

**Figure 4.**
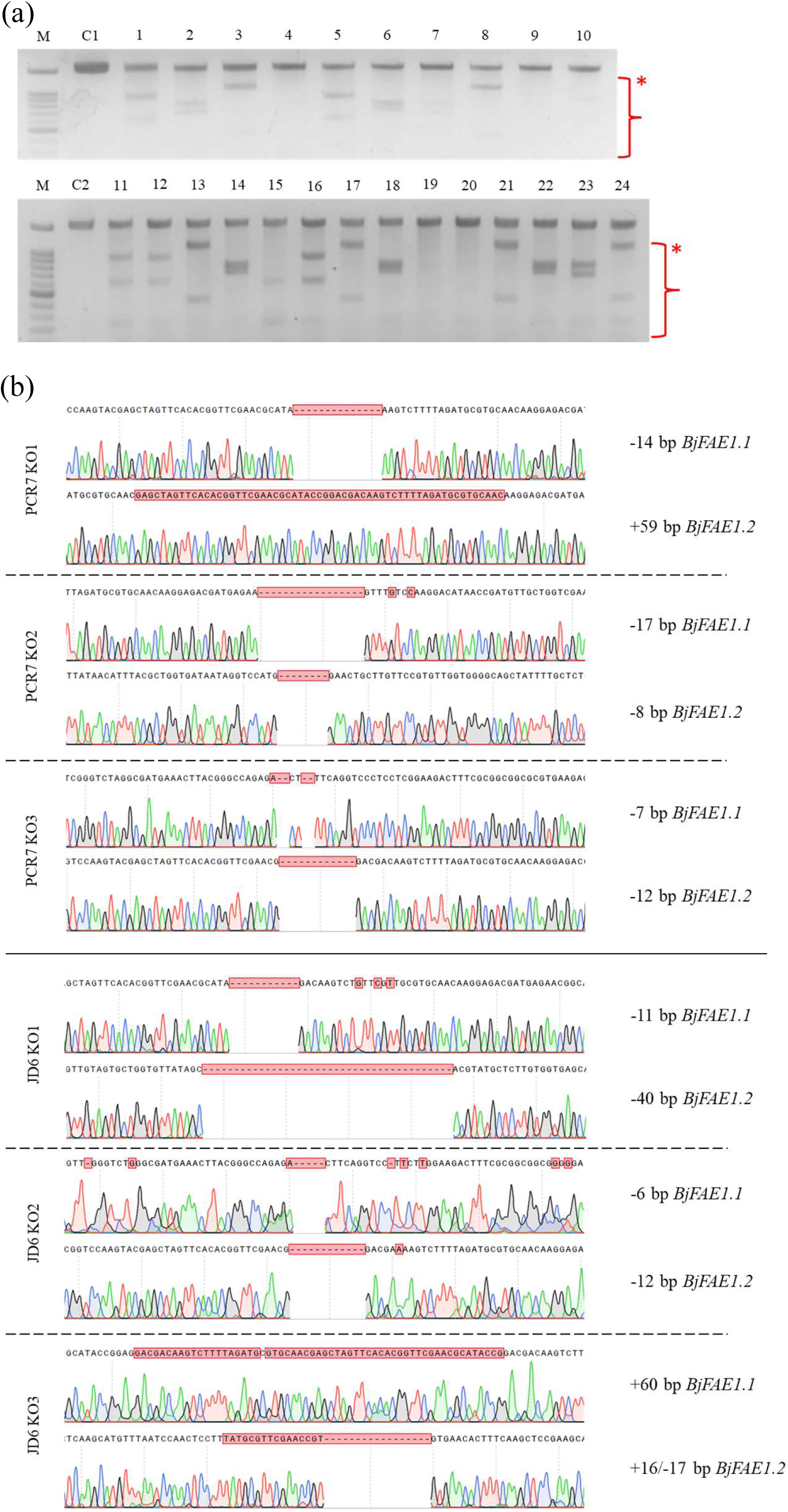
Molecular confirmation of biallelic *BjFAE1* mutations (e1e2) in T2 lines of two cultivars, PCR7 and JD6. (**a**) T7 endonuclease I (T7EI) mismatch detection assay to identify CRISPR-mediated indels in 24 representative T2 transgenic plants. Lanes 1-10: mutant PCR7 lines; 11-24: mutant JD6 lines; C1: untransformed control of PCR7; C2: untransformed control of JD6; M: 100 bp DNA ladder. Red asterisks mark T7EI-cleaved fragments. (**b**) Sanger sequencing confirming Cas9/sgRNA-induced indels in *BjFAE1* homeoalleles of two representative T2 plants of each cultivar.

### Biallelic *BjFAE1* knockout lines demonstrated drastic reduction of EA content in *B. juncea* seeds

To assess the impact of *BjFAE1* knockout on seed oil quality, we performed a comparative FA profile analysis of mature T2 generation seeds using GC-MS. Biallelic *FAE1* KO lines (mutant homeoalleles denoted as e1 and e2) of PCR7 cultivar exhibited a substantial reduction in seed EA content, from approximately 40% in wild-type plants to less than 0.5% (**Figure 5a**, **5b**). This remarkable decrease in C22:1 was accompanied by a compensatory increase in the levels of desirable FAs, including C18:1, C18:2, and C18:3 FAs from ∼18% to ∼30%, ∼14% to ∼26%, and ∼8% to ∼20%, respectively, in seed oil (**Figure 5a**, **5b**). In a similar trend, single allele mutation of *BjFAE1.1* (mutant lines denoted as e1) and *FAE1.2* (mutant lines denoted as e2) in PCR7 cultivar also resulted in the reduced C22:1 content to ∼one-third in e1 and ∼one-half in e2 in seed oil compared to the untransformed control PCR7 plants (**Table S7**).

**Figure 5.**
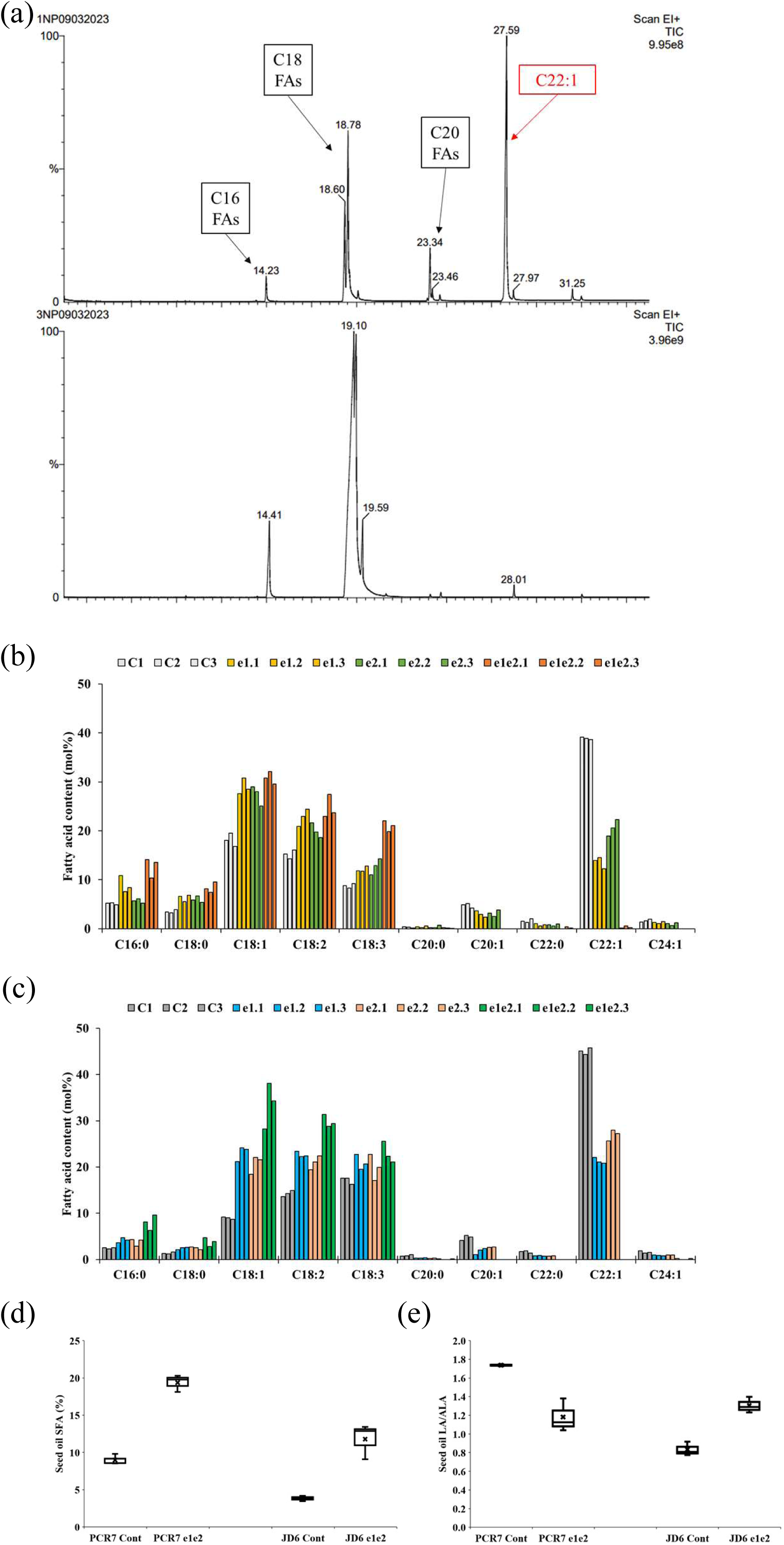
CRISPR/Cas9-mediated *BjFAE1* knockout alters seed FA composition in Indian mustard cultivars, PCR7 and JD6. (**a**) Representative GC-MS chromatograms of seed FA profile of untransformed control (upper panel) and CRISPR/Cas9-edited PCR7 cultivar (lower panel). (**b**) Seed FA compositions of T2 KO lines of PCR7 along with untransformed control, n=3 in each case. (**c**) Seed FA compositions of T2 KO lines of JD6 along with the untransformed control, n=3 in each case. (**d**) Seed oil SFA content (%) of e1e2 KO lines of two cultivars, n=3 in each case. (**e**) LA to ALA ratio of seed oil of e1e2 KO lines of two cultivars, n=3 in each case.

Comparable results were observed in seed FA profiles of JD6 lines. The e1e2 lines of JD6 cultivar showed a dramatic decrease in C22:1 from ∼45% to undetectable level, along with a corresponding increase in C18:1, C18:2, and C18:3 levels from ∼9% to ∼35%, ∼14% to ∼30%, and ∼17% to ∼24%, respectively, in seed oil (**Figure 5c**, **Figure S1**). Similar to PCR7 lines, JD6 e1 or e2 mutants also demonstrated a significant reduction in C22:1 content to ∼22% and ∼25%, respectively, in seed oil compared to the untransformed control JD6 plants (**Table S8**). Selected e1e2 lines of PCR7 and JD6 recorded higher seed oleic acid content of 32.05% and 38.08%, respectively, compared to ∼18% in PCR7 and <10% in JD6 control (**Figure 5b**, **5c**, **Table S7**, **S8**, **S9**). The average saturated FA (SFA) content of e1e2 lines increased from ∼9% to ∼ 21% and from ∼4% to ∼ 12% in case of PCR7 and JD6, respectively (**Figure 5d**, **Table S9**). Despite the substantial changes in overall FA composition, the linoleic to alpha-linolenic acid (LA/ALA) ratio in the e1e2 KO lines remained comparable to the untransformed control cultivars (**Figure 5e**, **Table S9**). Overall, these findings document that targeted knockout of *BjFAE1* genes effectively reduces the unwanted EA content while proportionally increasing the level of nutritionally desired oleic, linoleic, and linolenic acids in the resulting *B. juncea* seed oil.

### Investigation of phenotypic variation and other agronomically important traits in generated knockout lines

To assess the impact of *BjFAE1* KO on agronomic performance, transgenic e1, e2, e1e2 lines and untransformed control cultivars were grown to maturity in a contained greenhouse. Observations throughout the growth cycle, including flowering (**Figure S2**) and seed ripening revealed no significant developmental variations between the CRISPR modified and control plants. At maturity, e1e2 KO lines exhibited plant morphology (**Figure 6a**) including mature seed size (**Figure 6b**) comparable to untransformed controls. Detailed analyses of key agronomic traits such as plant height, branch number and inflorescence length, silique number and length, thousand-seed weight, and seed germination frequency demonstrated no significant alterations in the modified e1, e2, e1e2 lines compared to untransformed control cultivars (**Table 1**). Importantly, seed oil from KO lines demonstrated a substantial decrease in EA content without perturbing seed oil and protein levels compared to the untransformed control (**Figure 6c**, **Table S10**). Collectively, these findings strongly suggest that targeted *BjFAE1* knockout remarkably reduces EA level in seed oil of *B. juncea* without compromising seed germination, plant growth, and yield components.

**Figure 6.**
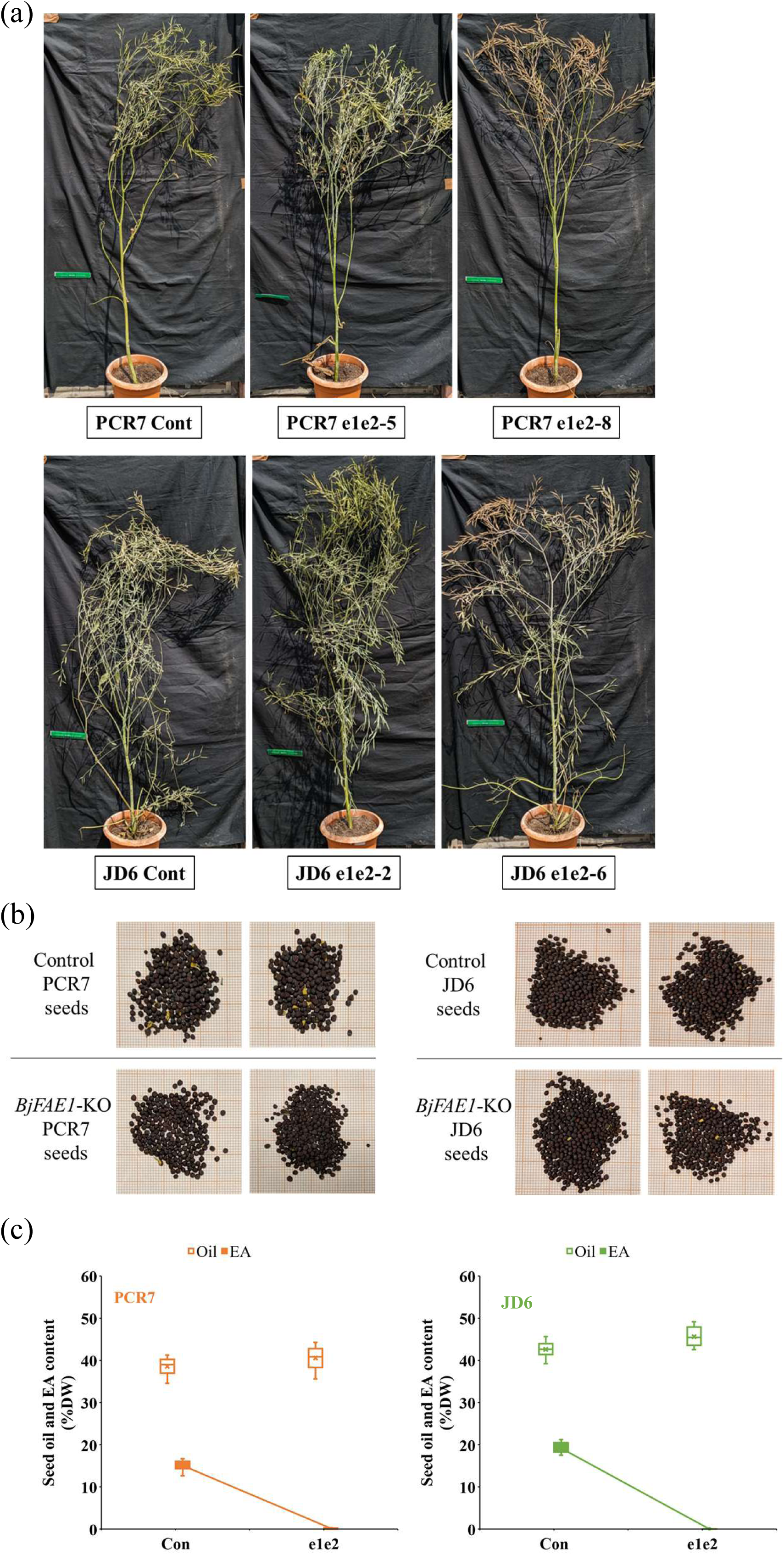
Agronomic trait investigation of e1, e2, e1e2 and untransformed control lines PCR7 and JD6 cultivars. (**a**) Plant morphology of untransformed control cultivars and two e1e2 lines of each cultivar during maturity. (**b**) Seed morphology of T2 generation of e1e2 KO lines along with their respective control cultivars. (**c**) Content (% DW) of seed oil and EA of e1e2 KO lines of two cultivars (n=5 in each case).

**Table 1.** Comparison of agronomic traits of PCR7 and JD6 cultivars with their monoallelic (e1 and e2) and biallelic (e1e2) KO lines. Values represent mean ± SE (n=10-12).

| Cultivar | Genotype | Plant height (cm) | Branches /plant | Inflorescence length (cm) | Siliqua /plant | Siliqua length (cm) | Seeds /siliqua | 1000 seed wt (g) | Germination frequency (%) |
| --- | --- | --- | --- | --- | --- | --- | --- | --- | --- |
| PCR7 | Control | 162.1±4.78 | 7.18±2.63 | 41.49±5.29 | 163.53±17.35 | 5.67±0.58 | 12.56±4.12 | 4.06±0.29 | 97.3±0.4 |
|  | e1 | 158.28±6.02 | 6.82±0.74 | 46.57±3.88 | 154.27±24.32 | 4.39±0.53 | 13.05±2.23 | 3.93±0.21 | 96.4±0.9 |
|  | e2 | 163.53±4.43 | 6.51±1.56 | 38.95±8.16 | 138.42±21.51 | 4.03±0.49 | 14.38±3.05 | 4.26±0.4 | 92.1±2.6 |
|  | e1e2 | 171.64±7.22 | 7.06±2.04 | 47.84±6.99 | 189.87±15.78 | 5.46±0.74 | 12.33±3.53 | 4.31±0.08 | 97.6±0.5 |
| JD6 | Control | 146.54±8.67 | 5.72±3.37 | 37.25±8.92 | 228.25±22.65 | 4.43±0.51 | 9.47±4.86 | 4.83±0.34 | 96.7±0.7 |
|  | e1 | 153.87±4.18 | 5.24±3.66 | 41.49±8.16 | 253.11±14.15 | 4.02±0.87 | 10.81±5.15 | 4.68±0.41 | 95.2±0.6 |
|  | e2 | 147.65±5.89 | 6.05±2.38 | 38.16±4.3 | 186.71±17.24 | 4.70±0.36 | 8.73±3.87 | 4.86±0.13 | 98.1±0.3 |
|  | e1e2 | 152.22±6.41 | 6.29±2.77 | 39.79±5.29 | 209.28±28.69 | 5.04±0.12 | 10.36±4.26 | 4.98±0.25 | 98.7±0.5 |

## Discussion

In plants, the initial steps of FA biosynthesis, leading to the formation of C18 chain length FAs, occur within the plastids (Ohlrogge and Browse, 1995). Subsequently, these C18 FAs are utilized by specialized fatty acid elongases like FAE1 on the endoplasmic reticulum, for further elongating C18-CoA chains into longer FA chains (Li-Beisson et al., 2010). The *FAE1* gene encoding ketoacyl-CoA synthase (KCS) plays the most critical role in EA biosynthesis (James et al., 1995). The gene was initially discovered in *Arabidopsis*, and subsequent researches have shown the presence of an expanded *FAE1* gene family in *Brassica* species, including *B. juncea* (Gupta et al., 2004). The EA (C22:1) biosynthesis pathway involves four enzymes, including KCS, 3-ketoacyl-CoA reductase (KCR), 3-hydroxyacyl-CoA dehydratase (HCD), and trans-2,3-enoyl-CoA reductase (ECR) for elongating C18:1 chain to C22:1 (Bach and Faure, 2010). Expression profile analysis, showing the positive correlation between *BjFAE1* transcript levels and seed EA content (**Figure 1c**), reaffirms the established role of FAE1 in EA accumulation. Consequently, targeting the *BjFAE1* gene has become a key strategy for developing LEA varieties. Traditionally, researchers obtained LEA mutants by either screening natural populations or by inducing random mutations using ionizing radiations (like gamma ray) or chemicals mutagens like ethyl methanesulfonate (Jambhulkar, 2007). More recently, RNA interference (RNAi) has been used to specifically suppress *BjFAE1* expression to reduce seed EA content (Sinha et al., 2007, Wang et al., 2022).

Our study demonstrates that CRISPR/Cas9-mediated knockout of *BjFAE1* homeoalleles (E1 and E2) effectively reduces EA content in seed oil of *B. juncea* seeds. This reduction in C22:1 is accompanied by a proportional increase in palmitic (C16:0), oleic (C18:1), linoleic (C18:2), and linolenic (C18:3) acids in seed oil. While the specific mechanisms underlying these changes in FA profile require further investigation, we can hypothesize that the absence of functional elongase enzyme disrupts the elongation of C18:1 substrate, which are the precursors of C22:1. Consequently, the carbon flux is rerouted in the FA biosynthesis pathway with accumulation of precursor FAs, leading to a proportional increase in the content of C16:0, C18:1, C18:2, and C18:3 **(Figure 5)**.

Although our *FAE1* knockout strategy demonstrated near zero seed EA content in biallelic KO e1e2 lines of both the cultivars used, the results (**Figure 5, 6d**) are consistent with previous studies employing RNAi-mediated silencing of *FAE1* in *B. napus* and *B. juncea* (Sinha et al., 2007, Wang et al., 2022), which also reported significant reductions in EA content alongside increases in oleic acid. This reinforces the crucial role of FAE1 enzyme in EA biosynthesis across *Brassica* species. However, CRISPR/Cas9 offers potentially greater precision and efficacy in achieving targeted gene disruption compared to RNAi, as documented by recent studies (Gan and Ling, 2022). CRISPR/Cas9 allows for direct modification of the target genomic DNA, potentially leading to stable gene knockout. Unlike RNAi, which exhibits variable degree of gene silencing in transgenic lines, CRISPR/Cas9-mediated gene knockout can create plants with the stable inheritance of LEA trait across generations.

Consistent with previous reports on CRISPR/Cas9-mediated genome editing in various crop species, the resulting indels in our study exhibited a wide range of sizes, from 60 base pair additions to deletions of 40 base pairs (**Figure 4b**). This diversity is in line with the expected variability of mutations induced by the non-homologous end-joining (NHEJ) repair mechanism that typically follows Cas9-mediated DNA cleavage. For example, studies in Arabidopsis, rice, tomato, maize, soybean, and wheat have reported indels ranging from few base pairs to few kilo base pairs (Gan and Ling, 2022). This broad spectrum of mutations highlights the flexibility of CRISPR/Cas9 as a tool for generating diverse genetic variations, which can be harnessed for crop improvement, in general.

Importantly, our work aligns with the growing body of research demonstrating the utility of CRISPR/Cas9 in *B. juncea* for diverse applications. Recent studies have successfully employed this technology to target the allergen Bra j I in *B. juncea*, reducing its allergenicity (Assou et al., 2022), and to modify the glucosinolate profile, creating low-seed, high-leaf glucosinolate oilseed mustard (Mann et al., 2022). The biallelic knockout of *BjFAE1* in our study resulted in a substantial decrease in EA content, reaching to an undetectable level in some T2 lines. This dramatic reduction coupled with a 1.8-fold (in PCR7) and 4.2-fold (in JD6) increase in oleic acid (**Figure 5**) brings those e1e2 lines much closer to the canola quality requirement of at least 50% oleic acid as per FCC standard. It is worthy to mention that the LA/ALA ratios in those biallelic KO lines fall within the recommended range (∼1-2:1) considered optimal for human health (Lands, 2008; Tu et al., 2010; Gibson et al., 2011). The higher proportion of oxidatively stable SFAs and oleic acid in the *e1e2* seed oil (**Table S9**) should also contribute to its longer shelf life. Previous studies along with our own findings, underscore the versatility and precision of CRISPR/Cas9 as a powerful tool for crop improvement in *B. juncea*.

An important limitation of our current approach is the presence of the Hyg^R^ gene expression cassette used as a selection marker during the CRISPR/Cas9 transformation process. While this marker effectively facilitated the identification of putative transgenic plant lines with edited *BjFAE1* gene, its presence falls under the regulatory purview of transgenic crops in many countries, potentially restricting the cultivation of these knockout lines. Future research efforts will focus on developing and employing marker-free strategies for CRISPR/Cas9-mediated *BjFAE1* knockout in *B. juncea*, which will significantly enhance regulatory compliance and broaden the potential adoption of these LEA varieties.

While the present study focused on the primary impacts on the nutritionally unwanted EA and the health beneficial oleic, linoleic, and linolenic acids; a complete metabolome could further explore non-verified changes in the spectrum of minor FAs and other lipid-derived molecules. Such changes could have implications for the edible oil’s oxidative stability or the production of industrially relevant oleochemicals. Additionally, future studies could investigate the long-term impact of *BjFAE1* knockout on plant stress tolerance or other physiological processes. Finally, evaluating the performance of these *BjFAE1* gene edited lines across diverse environmental conditions would be crucial for their potential deployment in field settings. These results advance the genetic resources available for breeding LEA *B. juncea* varieties and demonstrate the effectiveness of CRISPR/Cas9 for precise trait modification in this important oilseed crop. Development of low or zero erucic acid *B. juncea* cultivars present a critical step towards broader and safer consumer use of it as a nutritional edible oil.

## Experimental procedures

### Plant material and explant preparation

Seeds of five *B. juncea* (L.) cultivars viz. NPJ-112, NPJ-113, NPJ-124, PCR7, and JD6 were obtained from Bidhan Chandra Krishi Vishwavidyalaya, West Bengal, India. Mature seeds of *B. juncea* (PCR7 and JD6) were surface-sterilized using 1.0 % NaOCl with 0.1 % Tween and germinated on solid half diluted MS media (Duchefa Biochemie) in the dark. Hypocotyls and halved cotyledons from 6-day-old seedlings were used for genetic transformation.

### Estimation of seed lipid content and fatty acid profiling by gas chromatography

The seed lipid was extracted and measured by gravimetric methods (Bligh and Dyer, 1959) and are individual extracted samples are converted to fatty acid methyl esters (FAMEs) following the protocol reported in Bhattacharya et al., 2015. FAMEs extracted from each sample were analyzed using gas chromatography (GC) coupled with mass spectrometer (MS) using Elite-5MS column in CLARUS 690 GC/MS instrument (Perkin Elmer). The specific types of fatty acids were determined by matching them against a reference database (NIST library). To accurately measure the amount of EA present, a standard curve was created using known concentrations of methyl erucate (Merck) (**Table S1**).

For FAs other than EA, the peak areas were taken into consideration. The size of each peak in the chromatogram directly reflects the quantity of the corresponding FA in the sample. The chromatogram for each lipid sample was analyzed and the relative proportions of different FAs was calculated as a percentage of the total moles (mol%). These proportions were then plotted using a bar graph.

### Isolation of tissue-specific and stage-specific RNA and quantitative reverse transcription PCR (qRT-PCR)

Total RNA was extracted from different *B. juncea* tissues (root, shoot, leaf, flower, seed) at different growth stage using the RNeasy mini kit (QIAGEN) and subsequently qualitatively checked for quality in 1.5% agarose gel. Quantification of RNA concentration was performed in Nanodrop equipment (Thermo Scientific NanoDrop 2000). To remove any genomic DNA contamination from the extracted RNA samples, DNaseI (Sigma) treatment was performed using the kit following instructions as mentioned in the kit catalogue. Complementary DNA (cDNA) was prepared from total RNA extracted from different stages of tissues by using high-capacity cDNA reverse transcription kit (Applied Biosystems) as per kit manual instructions and used for real-time PCR experiments. *BjFAE1* transcript expression analysis was performed using Power SYBR Green Master Mix (Invitrogen) on StepOnePlus (Applied Biosystems) instrument following manufacturer’s protocol using elongation factor 1-alpha (*EF1A*) as reference (Dixit et al., 2019).

### Selection of gRNAs and preparation of CRISPR/Cas9 construct

Full-length coding DNA sequences (CDS) of *BjFAE1.1* and *BjFAE1.2* from the *B. juncea* genome were used as input sequences to design the 20 nt gRNAs using CRISPR-P v2.0 (Liu et al., 2017). Four single gRNAs independently targeting *BjFAE1* homeoalleles were selected viz., sgRNAs 1 – 4 based on their high on-target efficiency (maximum on-score) and low potential for off-target effects. Additionally, all selected sgRNAs contained the 3’ NGG protospacer adjacent motif (PAM) sequence, which is essential for the recognition and cleavage activity of *Streptococcus pyogenes* Cas9 on the target DNA. Selected *BjFAE1*-targeting gRNAs were synthesized as forward and reverse oligos and cloned at sgRNA1 (*Bsa*I-5′-GGTCTC(N1)/(N5)-3′) and sgRNA2 (*Esp*3I-5′-CGTCTC(N1)/(N5)-3′) oligo cloning positions of the dual guide accepter plasmid (from Prof. Guy C. Barker, University of Warwick, UK) for simultaneous targeting a pair of genomic regions. After transformation of *Escherichia coli* TOP10 competent cells (Thermo Fisher Scientific) with the individual recombinant plasmids, oligo assemblies at *Bsa*I site were verified by generating 590 bp amplicon from colony PCR with LI152 as forward primer and position I guide complementary as reverse primer (**Figure S3**) followed by sequence analysis of the amplicon. Similarly, *Esp*3I oligo clones were verified by generating 327 bp amplicon from colony PCR with guide 1 as forward primer and position II guide complementary as reverse primer (**Figure S3**), followed by sequence analysis of the amplicon. All the guides and primers used in this study listed in **Table S4** and **Table S11**. Two such chimeric constructs viz., pBjFAE13 and pBjFAE24 each harboring dual sgRNAs of different combinations were made (**Figure 2C**) and confirmed by sequencing.

### *Agrobacterium*-mediated *B. juncea* transformation, transformant selection and plant regeneration

The chimeric gene constructs, pBjFAE13 and pBjFAE24, were individually introduced into competent *Agrobacterium tumefaciens* strain LBA4404/VirGN54D (Fits et al., 2000) and positive clones are selected for *B. juncea* transformation **(Figure S4)**. Hypocotyl explants from 6-day-old PCR7 and JD6 seedlings were used for transformation. Putative transformants were selected using 30 mg/l of hygromycin (Duchefa Biochemie) and regenerated following the protocol reported by Bhattacharya et al. (2015) **(Figure S5)**.

### Isolation of genomic DNA and screening of mutant T0 plants

Genomic DNA (gDNA) was isolated from the young leaves for mutation analysis using the CTAB method (Doyle and Doyle, 1987). For High-Resolution Melting (HRM) screening, equal amount of gDNAs were pooled from untransformed control plant and sets of putative transformants. Approximately 150-200 bp of the genomic sequence flanking the target sites of each *BjFAE1* homeoalleles were amplified with the gene-specific primers (**Table S11**) using MeltDoctor HRM Master Mix (Applied Biosystems). HRM experiment were performed on StepOnePlus instrument (Applied Biosystems) following manufacturer’s protocol. Melting curves were generated by gradually increasing the temperature from 65 °C to 95 °C in 0.2 °C increments. The resulting melting curves were analyzed using HRM v3.0.1 (Applied Biosystems). PCR products generated using HRM primers and Q5 High-Fidelity DNA Polymerase (New England Biolabs) was gel-eluted and sequenced by Sanger sequencing. To identify mutations in the target genes, the sequence chromatogram files of each edited/mutant plant line were compared with those of the control non-edited plant using the ICE analysis tool version 3.0 (Synthego Corporation) (Conant et al., 2022).

### Confirmation of *FAE1* KO plants in T2 generation and off-target mutation detection

Harvested T1 seeds were germinated in the presence of the antibiotic hygromycin to select for the presence of *A. tumefaciens* T-DNA, confirming the successful transformation events. Plants were then transferred to greenhouse conditions till seed harvesting.

To identify CRISPR-mediated insertion-deletion (indel), T7 endonuclease I (T7EI) mismatch detection assay was performed. Briefly genomic DNA was extracted from hygromycin-resistant plants, as well as control plants and a region of interest encompassing the probable Cas9 cut site was amplified using specific primers (Table S11) using Q5 High-Fidelity DNA Polymerase (NEB). Equal amount of PCR products from putative transgenic plant and respective control plant were mixed and then denatured and reannealed, promoting the formation of heteroduplexes between wild-type DNA and mutant strands potentially containing indels. Heteroduplexes were treated with T7EI, which recognizes and cleaves mismatches (indel sites), while leaving wild-type DNA intact. Cleaved fragments were resolved and analyzed on an agarose gel. Plants exhibiting positive results in the T7E1 assay were selected for further analysis. Full-length *BjFAE1* sequences were cloned from these plants and subjected to Sanger sequencing to confirm the presence of indels. The edited and control sequences were aligned using the SnapGene software (GSL Biotech). Top two potential off-target sites were predicted using CRISPR-P2 (Liu et al., 2017) against each sgRNA. Total 8 sites were PCR amplified using primer pairs (Table S11) encompassing the off-target site sequenced from three e1e2 lines of each cultivar at T2.

### Determination of seed germination percentage

Seed germination rates were assessed for control, T0, and T1 transgenic plants, both in vitro and in greenhouse conditions. For in vitro germination, 20 seeds from each plant were randomly chosen 20 surface-sterilized seeds per plant were placed on solid half diluted MS media and kept in dark for 7 days at 25 °C. A seed was considered as germinated when the radicle and the hypocotyl had emerged within this period. For greenhouse germination experiments, 30 seeds from control, T0, and T1 transgenic plants were directly sown into 10-cm pots containing Soilrite mix (Keltech Energies) under a 14/10 h day-night cycle, at 25 °C. The germination was assessed after 14 days, once the cotyledons had emerged above growing medium.

### Investigation of agronomic traits

T1 and T2 mutant plants of e1, e2 and e1e2 lines and untransformed plants were grown in a greenhouse (16/8 h of light/dark at 22 °C) in 2023 and 2024, respectively. Various growth parameters such as plant height, branch number and inflorescence length, silique number and length, thousand-seed weight, and seed germination frequency of all the mutant lines and the control plants were measured throughout their development starting from seed germination to seed maturity stages.

## Acknowledgements

This research was funded by DBT (India) grant BT/IN/UK/PORI/03/AKP/2018-19 awarded to MKM and partially by BBSRC (UK) grant BB/R019819/1 awarded to GCB. NP acknowledges financial support from UGC (India). We thank Dr. Prabir Kumar Bhattacharyya (Associate Professor, Department of Genetics and Plant Breeding, Bidhan Chandra Krishi Vishwavidyalaya, India) for providing the *B. juncea* cultivar seeds used in the study. We also gratefully acknowledge the use of central facilities at UoW, UK and IIT Kharagpur, India, as well as the technical assistance of Dr. Luca Illing, Mr. Susamoy Sarkar, Mr. Bharat Dagra, and Mr. Nitai Giri.

## Author contributions

MKM and GCB planned the research work and designed the experiments. NP performed the experiments. NP, GCB, and MKM analyzed the data, discussed the results. NP wrote the manuscript with comments from all authors. All the authors read and approved the final article.

## Supporting information

**Figure S1** Representative GC-MS chromatograms of seed FA profile of untransformed control (upper panel) and e1e2 KO (lower panel) JD6 cultivar.

**Figure S2** Plant morphology of untransformed control plant (labeled as Cont) and two e1e2 lines of PCR7 and JD6 cultivar during flowering time.

**Figure S3** Preparation of pBjFAE13 and pBjFAE24 chimeric constructs.

**Figure S4** Selection of *A. tumefaciens* LBA4404 clones harboring either pBjFAE13 or pBjFAE24 KO construct.

**Figure S5** Schematic presentation of steps followed for *Agrobacterium*-mediated transformation of *B. juncea* hypocotyls with the KO constructs.

**Table S1** Estimation of seed oil content (%dry weight, %DW) and erucic acid content of five Indian mustard cultivars.

**Table S2** Seed fatty acid composition of five Indian mustard cultivars.

**Table S3** *BjFAE1* gene expression profile along with corresponding EA content in developing seed tissues of selected *B. juncea* cultivars.

**Table S4** Designed four single guide (sg) RNAs and their respective target sequence of *BjFAE1* coding sequence.

**Table S5** Analysis of CRISPR/Cas9-induced mutation in the *BjFAE1* homeoalleles in five representative *BjFAE1*-edited lines in the T0 generation of each cultivar.

**Table S6** Inheritance pattern of T-DNA in transgenic T1 plants of *B. juncea* cultivars PCR7 and JD6.

**Table S7** Seed fatty acid compositions of different T2 KO lines of PCR7 cultivar along with untransformed control.

**Table S8** Seed fatty acid compositions of different T2 KO lines of JD6 cultivar along with untransformed control.

**Table S9** Seed saturated fatty acid (SFA) content, oleic acid content and LA to ALA ratio e1e2 KO T2 lines with untransformed controls of PCR7 and JD6 cultivars.

**Table S10** Seed oil and protein content of e1, e2 and e1e2 KO T2 lines with untransformed controls of PCR7 and JD6 cultivars.

**Table S11** List of primers used in this study.

