## Supplementary figures for "Knockout of *fatty acid elongase1* homeoalleles in amphidiploid *Brassica juncea* leads to undetectable erucic acid in seed oil"

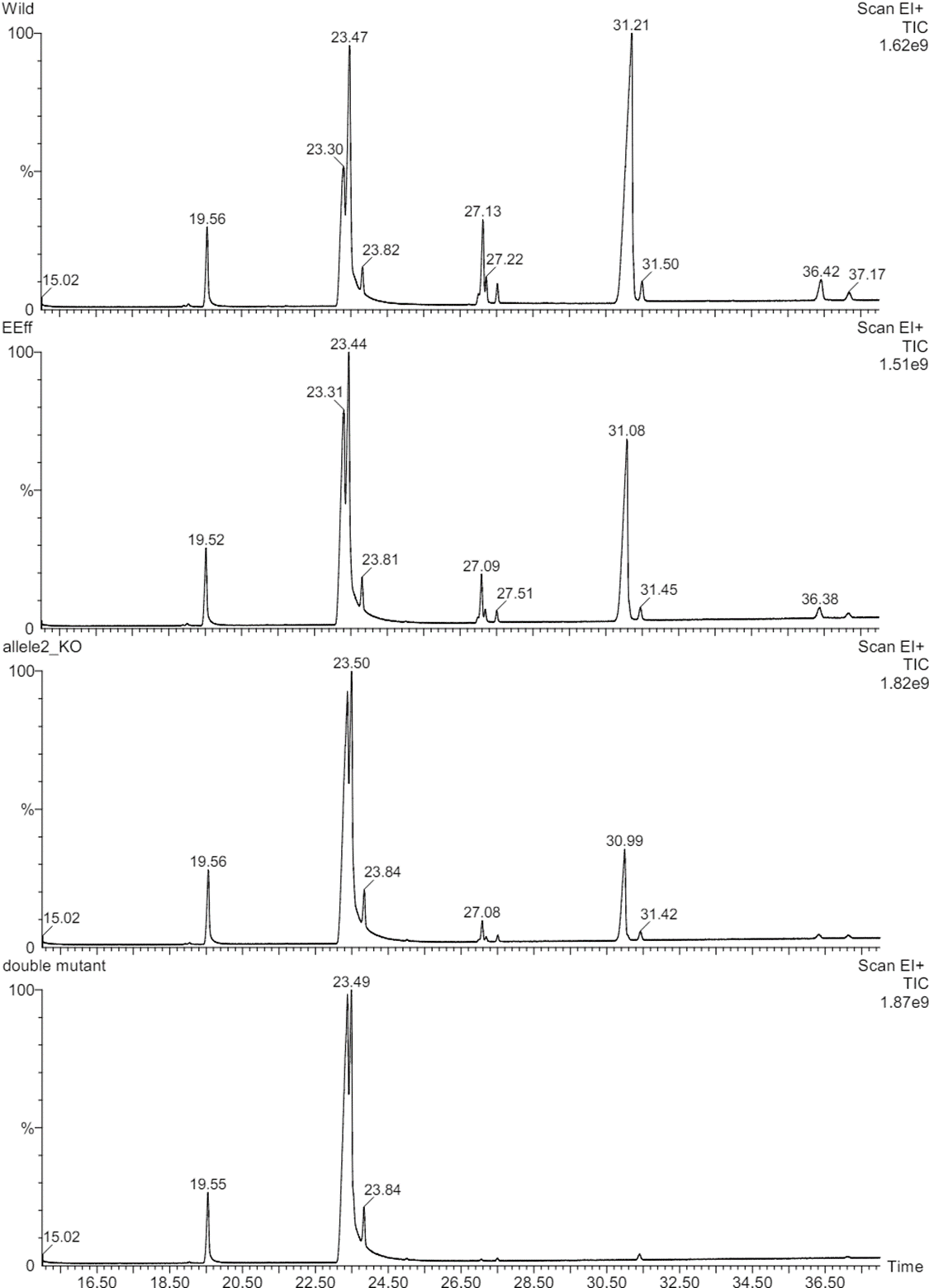

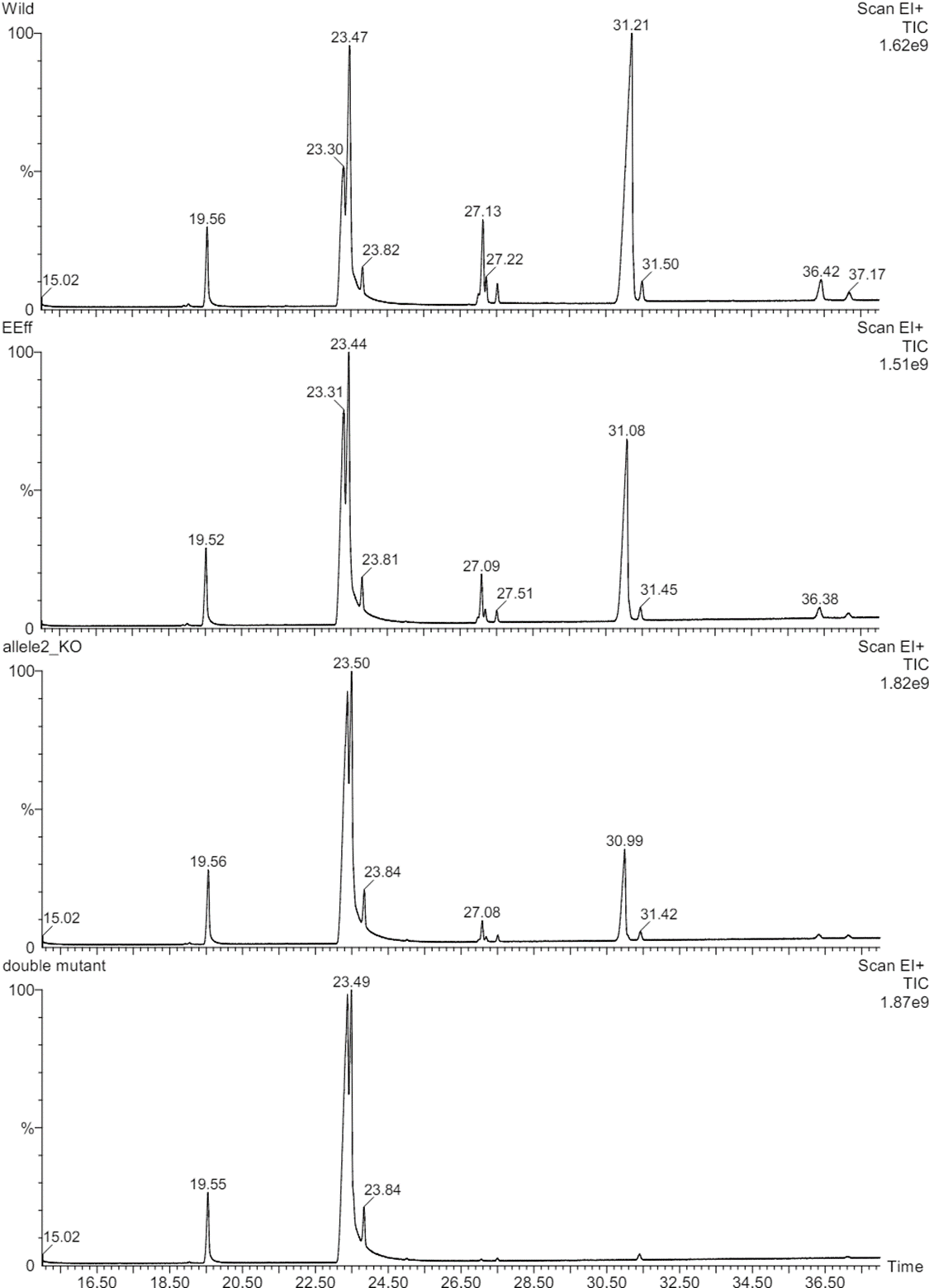

C22:1

**Figure S1** Representative GC-MS chromatograms of seed fatty acid profile of untransformed control (upper panel) and e1e2 KO (lower panel) JD6 cultivar.

(a)
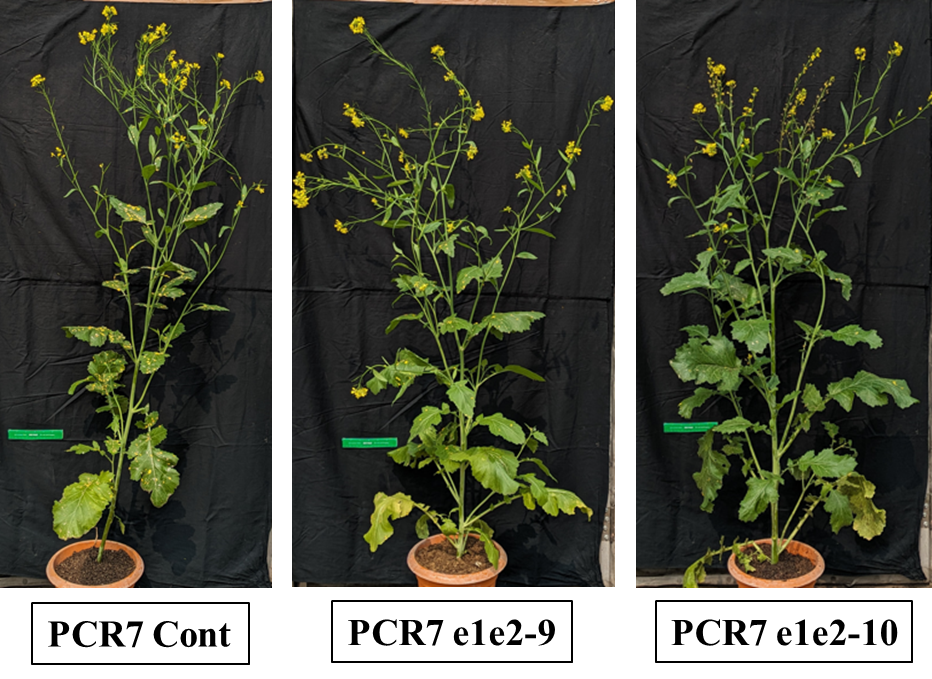

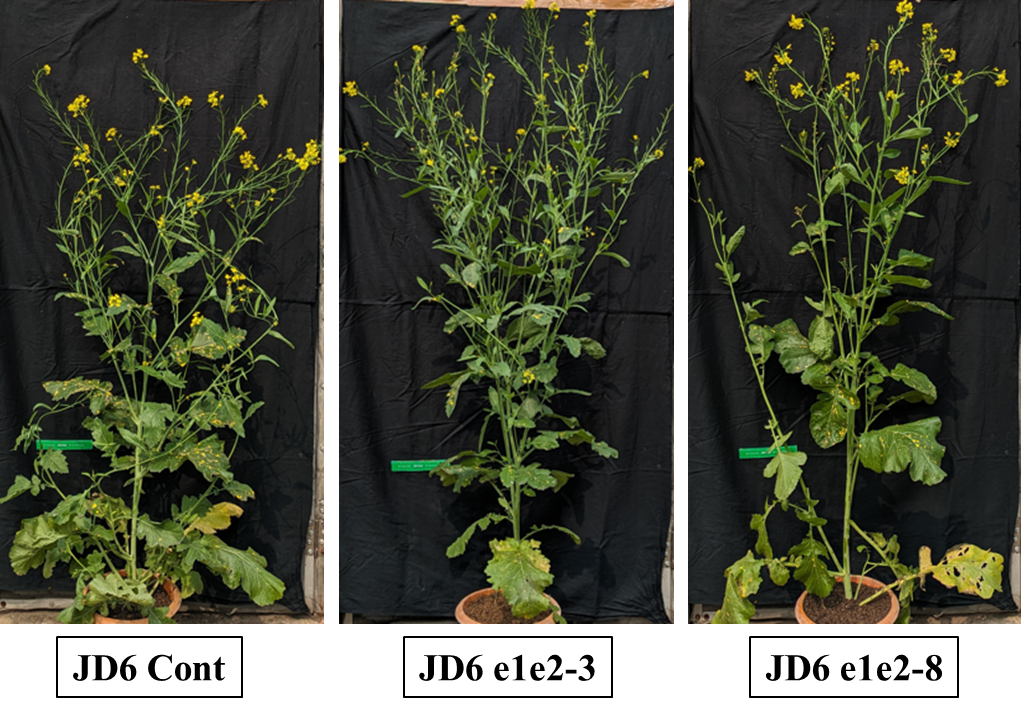
(b)

**Figure S2** Plant morphology of untransformed control plant (labeled as Cont) and two e1e2 lines of **(a)** PCR7 cultivar and **(b)** JD6 cultivar during flowering time. Observation shows there is no significant alternation in plant morphology between unedited control and KO lines.

(a)

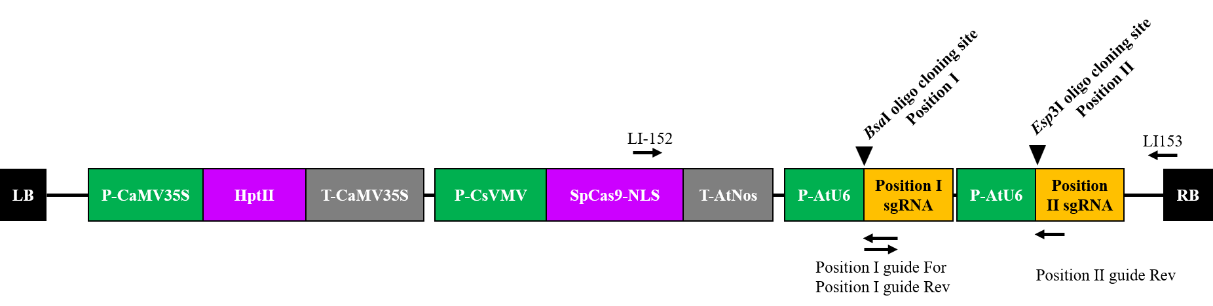

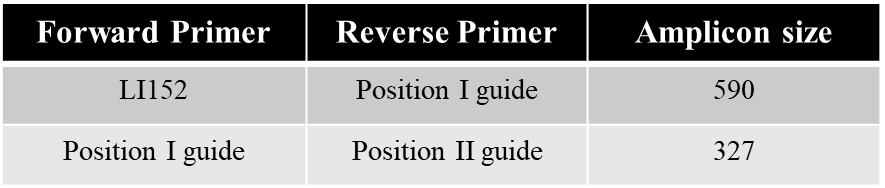

(b)

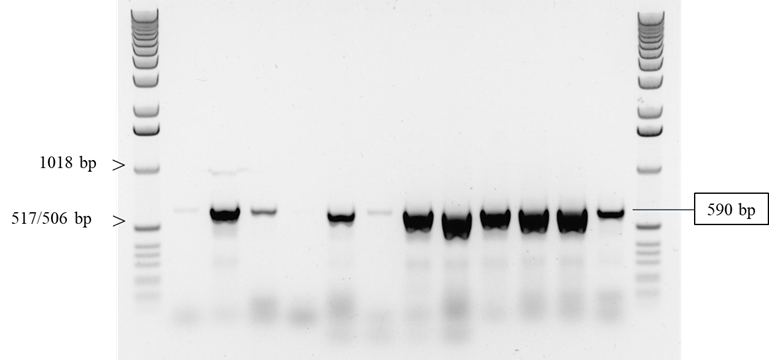

(c)

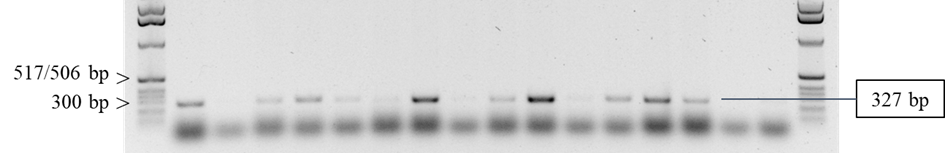

**Figure S3** Preparation of pBjFAE13 and pBjFAE24 chimeric constructs. **(a)** Schematic representation of dual accepter binary plasmid ready to receive oligos at position I and position II via *Bsa*I and *Esp*3I restriction enzyme sites, respectively, in order to allow target specific mutagenesis. Shown are the PCR primer locations used for positive clone identification before Sanger sequencing. **(b)** Agarose gel picture showing the amplicons obtained after colony PCR with LI152 forward and position I guide reverse primers for verification of oligo cloning at *Bsa*I site. **(c)** Agarose gel picture showing the amplicons obtained after colony PCR with position I guide as forward and position II guide as reverse primers for verification of oligo cloning at *Esp*3I site.

(a)

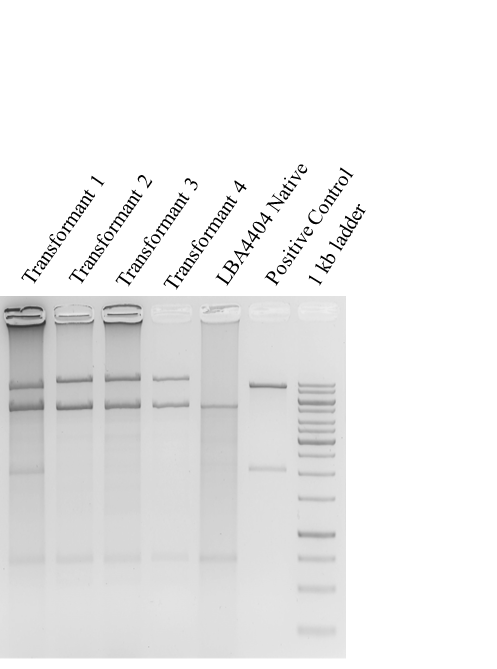

(b)

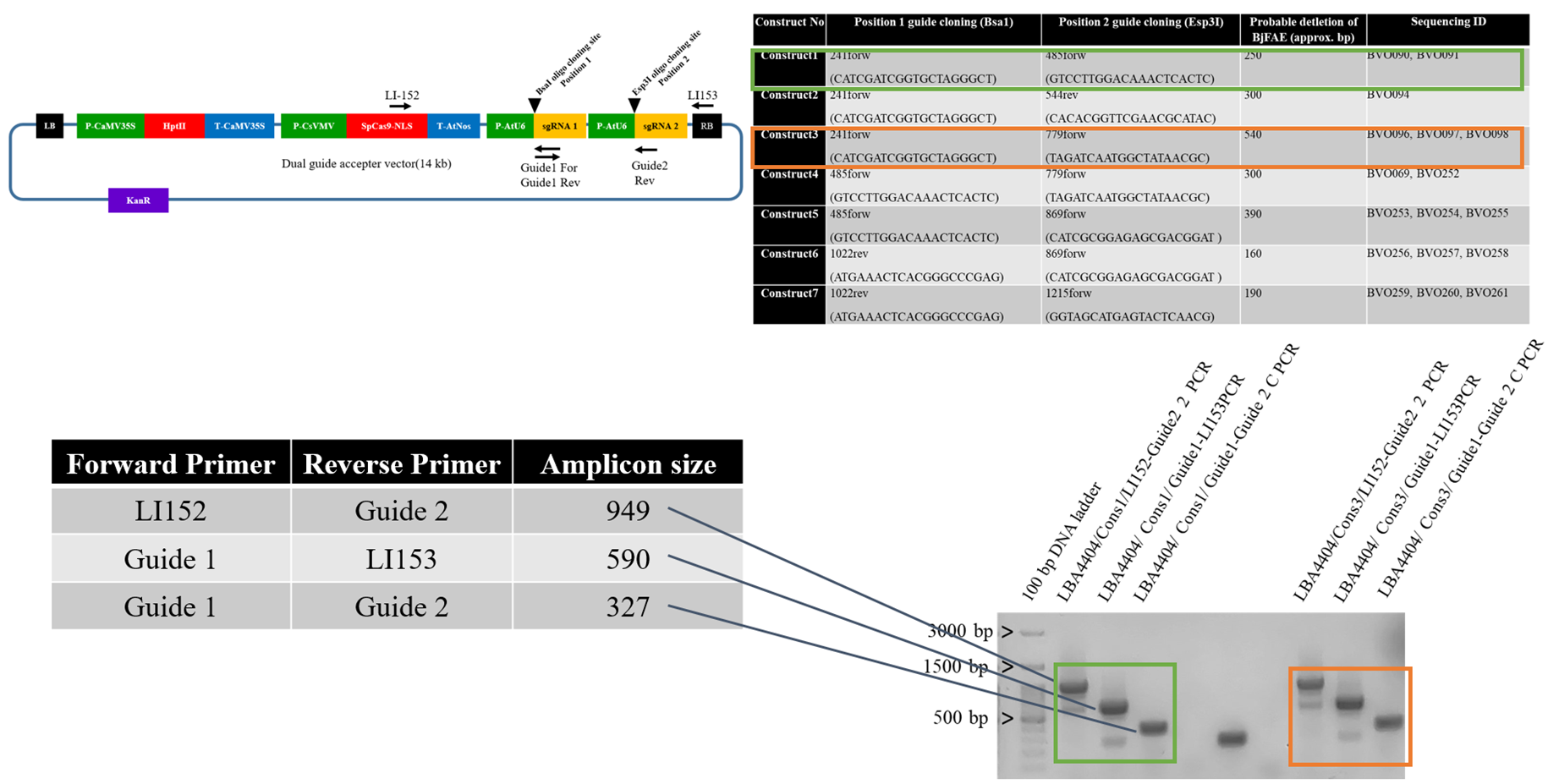

**Figure S4** Selection of *A. tumefaciens* LBA4404 clones harboring either pBjFAE13 or pBjFAE24 KO construct. **(a)** Screening of positive LBA4404 transformants carrying pBjFAE13 (Transformant 1 and Transformant 2) and pBjFAE24 (Transformant 3 and Transformant 4). Agarose gel showing the undigested plasmid DNAs isolated from the respective LBA4404 transformants along with the pBjFAE24 DNA isolated from the *E. coli* TOP10 clone as positive control. **(b)** Verification of LBA4404 positive clones by PCR screening using different primer pairs listed for pBjFAE13 (highlighted by green in agarose gel picture) and pBjFAE24 (highlighted by orange in agarose gel picture).

(a)

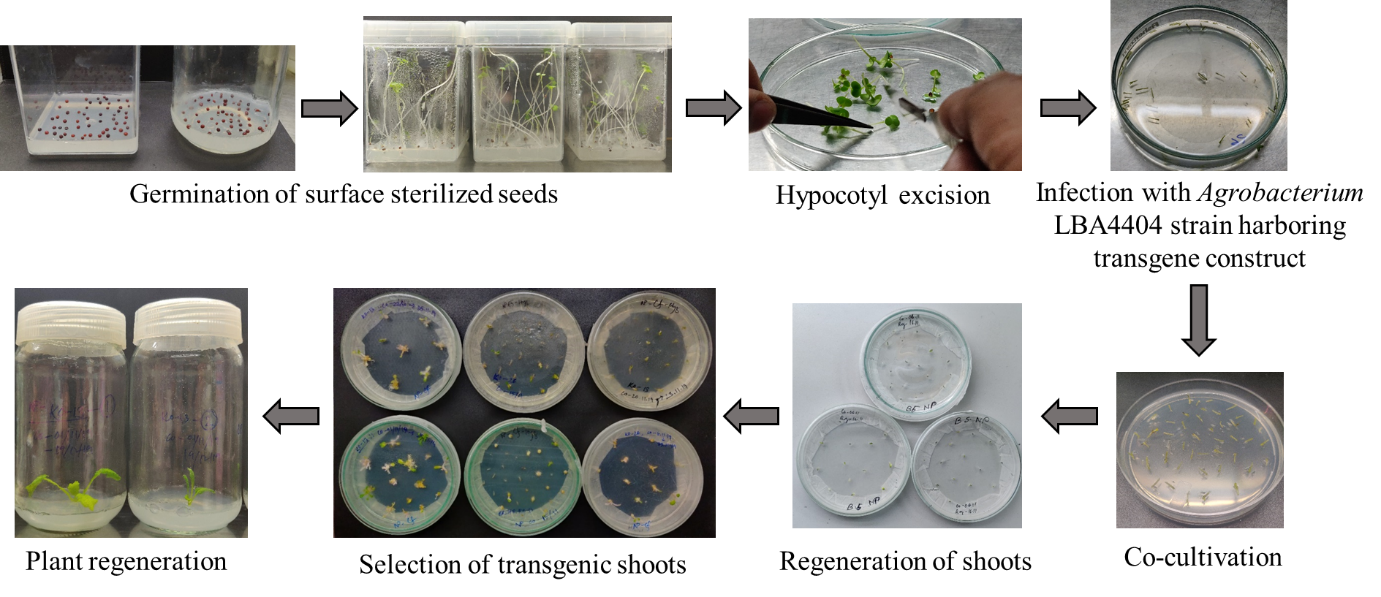

**Figure S5** Schematic presentation of steps followed for *Agrobacterium*-mediated transformation of *B. juncea* hypocotyls with the KO constructs. Shown are the pictures for PCR7 cultivar transformed with pBjFAE13construct
